# Dicamba drift alters plant-herbivore interactions at the agro-ecological interface

**DOI:** 10.1101/2021.07.13.452219

**Authors:** Nia M Johnson, Regina S Baucom

**Affiliations:** Ecology and Evolutionary Biology Department, 4034 Biological Sciences Building, University of Michigan, Ann Arbor, MI 48109

**Keywords:** agroecology, eco-evolution, plant-insect interactions, herbicide drift, weed adaptations

## Abstract

Natural populations evolve in response to biotic and abiotic changes in their environment, which shape species interactions and ecosystem dynamics. Agricultural systems can introduce novel conditions via herbicide exposure to non-crop habitats in surrounding fields. While herbicide drift is known to produce a variety of toxic effects in plants, little is known about its impact on non-target wildlife species interactions. In a two-year study, we investigated the impact of herbicide drift on plant-herbivore interactions with common weed velvetleaf (*Abutlion theophrasti*) as the focal species. The findings reveal a significant increase in the phloem feeding silverleaf whitefly (*Bermisia tabaci*) abundance on plants exposed to herbicide at drift rates of 0.5% and 1% of the field dose. We also identified a significant phenotypic tradeoff between whitefly resistance and herbicide resistance in addition to whitefly resistance and relative growth rate in the presence of dicamba drift after increasing the populations grown in year two. In a follow-up greenhouse study, we found evidence that dicamba drift at 0.5% of the field dose significantly increased average chlorophyll content (mg/cm^2^) along with a positive correlation between whitefly abundance and chlorophyll content. Overall, these findings suggest herbicide exposure to non-target communities can significantly alter herbivore populations, potentially impacting biodiversity and community dynamics of weed populations found at the agro-ecological interface.

## Introduction

Biotic and abiotic interactions influence species assemblage and evolution throughout ecosystems (Thompson 1999, Klanderud et al. 2015). Often a source of atypical abiotic factors, agricultural practices are largely dependent on pesticide use for increased crop yield and labor efficiency (Gianessi 2013). Despite the economic benefits of herbicides for weed control, however, their use can lead to unintentional impacts on non-target wildlife if the herbicide ‘drifts’ or migrates outside of the target area during or after the initial application (Freemark and Boutin 1995, Carlsen et al. 2006a, Carlsen et al. 2006b). Though herbicide drift is known to produce a variety of toxic effects on non-target vegetation (Marrs et al. 1991, Fletcher et al. 1996, Marrs and Frost 1997, Gove et al. 2007, Boutin et al. 2014, Cederlund 2017), and can lead to significant plant phenotypic and compositional changes (Iriart, Baucom, and Ashman 2020), few studies have evaluated how herbicide drift may disrupt or influence the interactions between plants and other community members such as pollinators and herbivores.

Weeds and other non-crop plants found at the edges of crops serve as important reservoirs of insect biodiversity (Egan and Mortensen 2012), which provides crucial ecosystem services for agriculture such as pollination and pest control (Daily 1997). The plant-insect interactions that occur at the agro-ecological interface are key determinants for the movement of energy and nutrients as well as drivers of ecological and evolutionary dynamics (Ehrlich and Raven 1964, Futuyma and Agrawal 2009). However, weed communities at the agro-ecological interface can be exposed to herbicide drift rates between 0.1% and 5% of the normal field application rate of herbicide (Cessna et al. 2005), and we currently understand very little about how this drift may influence ecological interactions and potentially lead to subsequent evolutionary responses. For example, does herbicide drift change herbivore abundance or damage, and/or lead to altered patterns of plant investment in resistance to either herbicide or herbivore damage?

One herbicide known for non-target damage is dicamba (3,6-dichloro-2-methoxybenzoic acid) (Jones et al. 2019). Dicamba is a broadleaf selective synthetic auxin that is on an upwards trajectory in agriculture due to a substantial increase in the adoption of dicamba-tolerant crops (USDA-ERA, 2019), which is evident by an estimated 600% increase of use in the US between 2014 and 2019 (Baker, 2021). Dicamba is absorbed by plant leaves and mimics the deformative and growth-altering effects caused by overdoses of the natural plant auxin, indole-3-acetic acid (IAA) (Grossman 2010). Dicamba drift can cause a shift in plant and arthropod diversity (Egan et l. 2014), delay the onset of flowering, reduce the number of flowers, and even reduce the amount of pollinator visits (Bohneblust et al. 2016). However, we have very little information on how dicamba drift may impact plant-herbivore interactions. A broad expectation is that exposure to a synthetic auxin, such as dicamba, could alter plant allocation towards growth and defense, and potentially lead to increased vulnerability to herbivores, in line with growth-defense tradeoff framework (Coley, Bryant, Chapin 1985). Recent work has shown that exposure to drift rates of herbicides like dicamba can lead to the evolution of reduced herbicide sensitivity (Vieria et al. 2020) and recurrent selection of low dose herbicides can lead to non-target site resistance (Busi et al. 2013), however, there is little understanding of how this process may influence selection for allocation herbivory defense.

Here we examined the impact of dicamba drift on plant herbivory, dominated by naturally occurring populations of the phloem feeding silverleaf whitefly (*Bermisia tabaci*) in the field, using the crop weed velvetleaf, *Abutilon theophrasti*. First introduced to the United States from Asia in the late 17th century, this annual species is now one of the most common broad-leaved species in and around corn and soybean agricultural fields located in Canada, Europe, and the United States (Spencer 1984). While past work in *Amaranthus hybridus* has shown that lines with evolved herbicide resistance (defined by survival given application of herbicide at the recommended field dose) can exhibit reduced herbivory resistance depending on light availability (Gassman 2005), there is a striking lack of information on how such a phenotypic tradeoff evolves when populations are exposed to drift rates under natural field conditions. We performed field experiments to address this gap in our knowledge, and specifically asked the following questions: Does the application of dicamba drift influence herbivory in the common weed *Abutilon theophrasti* (velvetleaf)? Does the amount of herbicide damage, which reflects resistance to herbicide, influence the amount of herbivory, which reflects resistance to herbivory? (*i*.*e*., is there a trade-off between these two forms of defense in the presence of dicamba drift)? Is either herbicide or herbivore resistance negatively correlated with relative growth rate, indicating a growth-defense tradeoff? Is there evidence of selection acting directly on herbicide resistance, herbivory resistance, or relative growth rate or evidence of correlative selection acting on the interactions of these focal traits? Does herbicide drift impact leaf chlorophyll levels, and if so, does that correlate with whitefly abundance?

## Methods

### Field Experimental Design

We performed two consecutive field experiments to characterize the extent of herbicide drift damage and herbivory response in *A. theophrasti*. In the first experiment, we planted replicate seeds from 23 maternal lines originally sampled from a single population (Dexter, MI) in the summer of 2018. We increased the number of populations from 1 to 8 for a total of 50 maternal lines in the second field experiment (SFig 1), including the 23 maternal lines from the originally sampled population and 2-5 maternal lines from the subsequently sampled populations. All seeds were collected in the Fall prior to each summer in and around soybean fields located in Dexter, MI.

In the first field experiment (2018) we scarified and planted 276 total seeds in a randomized block design (23 maternal lines × 3 treatments × 2 replications × 2 blocks) with each treatment randomized within each block, and two replicate plants per maternal line present within each treatment/block combination. Our three experimental treatments were two levels of a drift exposure (1% and 0.5% of the recommended field dose of dicamba (3,6-dichloro-2-methoxybenzoic acid) (2.8 g ae/ ha dicamba, 5.6 g ae/ha dicamba) and water as the control. For the second field experiment (2019), we scarified and planted 450 total seeds in a randomized block design with the same treatments as the previous field experiment (50 maternal lines × 3 treatments × 3 blocks). Treatments were randomized in each block, and one replicate of each maternal line was present within each treatment/block combination. We recorded leaf count, height, and largest leaf width for all plants biweekly in both experiments. We recorded flower count weekly and sampled all seeds at the end of the season for an estimate of fitness. Five weeks after seed germination, when the average plant height was 11 cm tall, we applied 1% and 0.5% dicamba to plants in each respective treatment with a hand-held pressurized prayer for roughly 3 seconds which ensured adequate coverage of the entire plant.

#### Herbicide Damage

We assessed herbicide damage two weeks following herbicide application by recording the proportion of leaves exhibiting visual deformation (curling or cupping), and proportion of the plant damage was determined by evaluating the number of leaves with a cupping or curling pattern divided by the total number of leaves.

#### Herbivory Damage

To estimate resistance to herbivory in 2018, we assessed both physical damage from chewing insects and the abundance of phloem-feeding insects (dominated by silverleaf whiteflies). Because there was little evidence of chewing insects, only whitefly abundance was assessed in 2019. We assessed chewing herbivory damage five weeks after herbicide application using the imaging software, Fiji ImageJ version 3.0 (Schindelin et al. 2012). To do this, we collected 3 leaves at random from each plant and scanned them using the Canon CanoScan L110. To obtain the total surface area, we converted each image to binary format and used the particle analysis tool which quantifies total pixel number into centimeters. We then used a macro plug-in to place a grid with 0.1cm^2^ spacing over each image to estimate the amount of chewing herbivore damage per leaf.

Visual estimates of whitefly abundance were sampled in both field experiments by selecting 10 leaves at random per plant. Because most whiteflies feed and oviposit at the same location (Van Lenteren and Noldus 1990), estimates of percentage of leaf area covered by whitefly oviposits were captured by visually partitioning each leaf into quarters and evaluated for each section, as suggested in Johnson et al 2015. We estimated whitefly abundance on a scale from 0 – 5 (adapted from Banks 1954). A score of 0 meant no whitefly oviposits were present, 1 was 1 - 20% leaf coverage by whitefly oviposits, two was between 21 – 40 % leaf coverage, three was 41 - 60 % leaf coverage, four representing 61 - 80% leaf coverage, and five representing 81 - 100% leaf coverage.

#### Photosynthetic carbon dioxide response

To determine if there was a link between herbicide resistance and photosynthetic response to dicamba drift, we estimated photosynthetic efficiency by examining variation in photosynthetic carbon dioxide response curves (A-Ci Curve) using the Li-Cor 6400 portable photosynthesis system (Open System Vers, 4.0, Li-Cor, Inc., Lincoln NE) in 2018. During gas exchange measurements, we maintained cuvette conditions at a photosynthetic photon flux density (PPFD) of 1500 μmol/ m^2^s, air flow rate at 500 μmol/s, and leaf chamber block temperature being 30°C to match ambient air temperature. In order to measure the gas exchange, we first set the CO_2_ concentration reference at 400 μmol/mol, and we maintained the leaf under such conditions for ten minutes for adaptation and stabilization of leaf photosynthesis. We set a loop of changing reference CO_2_ concentration at 400, 300, 200, 100, and 50μmol/mol. We controlled CO_2_ concentration in the cuvette with a CO_2_ mixer across this series and measurements were recorded after equilibration to a steady state. We then set another loop of changing reference CO_2_ concentration at 400, 600, 800, 1000, 1300, and 1500 μmol/mol and again logged at each iteration. Between each group of measurements, we set the reference CO_2_ concentration back to 400 μmol/mol for 5 minutes. These measurements were made four to five weeks after herbicide application, within a ten-day period. We recorded these data on a leaf area of 2 cm^2^, from 2 leaves on each of three plants from each treatment. For plants growing in drift treatments, we included one leaf that developed prior to herbicide application (Leaf 1), and one leaf that developed after herbicide application which as a result exhibited a deformation in shape (Leaf 2). The purpose of taking measurements on leaves with different times of development was to investigate if potential herbicide sequestration impacts photosynthetic efficiency of leaves exhibiting damage, potentially impacting herbivore resistance mechanisms for the entire plant. Measurements for each leaf per plant were taken on the same day.

### Greenhouse Experimental Design

With the goals of understanding treatment effects on whitefly abundance (*i*.*e*. impacts on host-plant selection), we conducted a greenhouse experiment in the summer of 2021. We planted 6 replicates of 20 maternal lines randomly selected from 5 of the populations used in the 2019 field experiment totaling 240 seeds (20 maternal lines × 6 experimental replicate seeds × 2 treatments). Half of the plants were directly treated with dicamba at 0.5% of the field dose, while the other half was treated with water 4 weeks following planting. In preparation for herbicide application, plants were placed in rows outside of the greenhouse in order to prevent herbicide drift. We subsequently applied the herbicide solution in the same manner as in the field experiments and relocated the plants back into the greenhouse once they had visually dried. We elected to treat plants with the herbicide one week prior to the field experiment due to observed accelerated growth in the greenhouse. Because whiteflies are common pests in many greenhouses, and the origin of the whitefly population in the field had previously been connected to crops transplanted from the greenhouse in plots adjacent to our own, we elected to allow whiteflies to naturally migrate to velvetleaf plants without manipulation or introduction.

#### Chlorophyll Content

We estimated whitefly abundance with the same methodology as above. Since whitefly host selection has previously been linked to leaf wavelength emissions (Mound 1962), using the atLEAF Plus Digital Chlorophyll Meter, we measured the chlorophyll levels (mg/cm^2^) of 3 leaves per plant chosen at random and calculated the average for each plant.

## Data Analysis

### Herbivory Response to Drift

We conducted all statistical analyses in R studio (version 3.4.1, R Development Core Team). For each type of herbivory measurement in the field experiments (chewing damage and whitefly abundance), we performed analysis of variance to determine whether herbivory resistance differed in response to dicamba application for each year separately. Prior to performing ANOVAs, we transformed response variables using Tukey’s Ladder of Powers in the rcompanion package (Mangiafico 2016) to correct for non-normality of residuals. In 2018 (n = 118), we fit the following mixed linear model using lmer function of the lmer4 package (Bates et al. 2011):

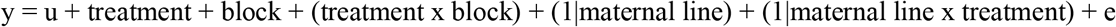

where, y, the response variable, is chewing damage or whitefly abundance, u is the intercept or mean of the model, treatment and block are fixed-effect terms, maternal line and maternal by treatment are random effect terms. In 2019 (n = 240), we nested maternal line into population and added population as a random effect to the existing model: y = u + treatment + block + (treatment x block) + (1|population) + (1|population: maternal line) + (1|population: maternal line) x treatment + e. We determined the significance of the predictor variables using *F*-statistics for the fixed effects with Kenward-Roger to estimate the degrees of freedom and used a log-likelihood ratio test to estimate χ^2^ for the random effect. We then used a Welch two sample t-test to determine if there were significant differences between the two dicamba drift treatments.

### Plant Performance in Response to Drift

To determine how drift treatments influenced plant performance in the field, we ran separate mixed linear models as described above by year. In 2018, we fit separate models for each plant trait (height, leaf count, leaf width, and flower number): y = u + treatment + block + (treatment x block) + (1|maternal line) + (1|maternal line x treatment) + e; where each trait was the response variable, block, treatment, and block by treatment interactions were fixed effects, and maternal line, and maternal by treatment interactions were random effects. In 2019, we nested maternal line into population and added population as random effects to the existing model: y = u + treatment + block + (treatment x block) + (1|population: maternal line) + (1|population: maternal line) x treatment + (1|population) + (1|population) x treatment + e. We transformed each trait again using Tukey’s Ladder of Powers to meet the assumptions of normality. We performed separate analyses for each year because preliminary analysis revealed a significant difference in plant growth between years.

Additionally, photosynthetic carbon dioxide response means were analyzed for 2018 using a Kruskal-Wallis test with treatment as the independent variable. We performed exact Wilcoxon rank sum tests on photosynthetic carbon dioxide response means of individuals grown in each drift treatment separately to determine if time of leaf development impacted photosynthetic carbon dioxide response among leaves with and without deformation by including leaf developmental time as the independent variable.

### Genetic Variation for Herbicide Resistance

We examined herbicide resistance for each year separately and were explicitly interested in determining if there was evidence for genetic variation for resistance to herbicide drift *via* a population or maternal line by treatment interaction. We fit the following separate mixed linear models for proportion of plant damaged from herbicide: y = u + treatment + block + (treatment x block) + (1|population: maternal line) + (1|population: maternal line) x treatment + (1|population) + (1|population) x treatment + e; where each type of resistance was the response variable, block, treatment, and block by treatment interactions were fixed effects, and maternal line, maternal by treatment interactions, population, and population by treatment interactions were random effects. We determined if there was evidence of genetic variation for herbicide resistance and whitefly resistance by performing log-likelihood ratio tests to estimate χ^2^ for maternal, maternal line by treatment, population, and population by treatment interactions for each resistance type. Finally, because we are interested in potential growth-defense trade-offs, we used a similar mixed linear model to examine genetic variation in relative growth rate.

### Operational Definition of Resistance

We define herbicide resistance operationally as 1 minus the proportion of leaves exhibiting damage from the herbicide, which was observed as yellowing and curling/deformation of leaves. Because there was limited damage from chewing herbivores, we elected to focus on phloem-feeding herbivory, dominated by whiteflies. Whitefly resistance was defined as 1 minus the amount of whitefly larvae coverage per individual (as described previously).

### Phenotypic Correlations

In order to test for correlations between herbicide resistance and whitefly resistance (defined operationally as above), we performed Pearson’s correlation test. We examined correlations for each experimental year separately.

### Phenotypic Selection

In accordance with the Lande–Arnold approach (e.g., Arnold et al. 2001; Arnold 2003; Hereford et al. 2004), we used phenotypic measurements to quantify natural selection on whitefly resistance, herbicide resistance and relative growth rate. Here, we report selection gradients rather than differentials to understand whether direct selection is acting on individual traits by controlling for any indirect selection present. Relative fitness was calculated as the final seed count per individual divided by mean seed count for each treatment per year (2018: n = 118; 2019: n = 240). We estimated linear (directional) selection gradients (*β*) within drift environments (0.5% and 1% the field dose of dicamba) by performing multiple linear regressions of relative fitness on whitefly resistance, herbicide resistance, relative growth rate (calculated as change in leaf count plus change in height divided by the number of weeks spent growing) and their interactions separately for each year. We also estimated nonlinear selection gradients (*γ*) in a full model that included linear terms, quadratic terms, and the cross-product terms of focal traits. Quadratic regression coefficients were doubled to estimate nonlinear selection gradients. Nonlinear selection gradients examined the potential for selection on phenotypic variance of a trait (quadratic selection) or phenotypic covariance (correlated selection) between focal traits, as there is evidence of relationships between plant stress, defense, and growth (Huot et al. 2014, Zust and Agrawal 2017). For all selection analysis we mean standardized focal traits (i.e., subtracted the mean and divided by the standard deviation) and used untransformed values of relative fitness. Furthermore, we estimated linear and nonlinear selection on whitefly resistance in the absence of herbicide drift in order to determine whether herbicide altered the pattern of selection. We compared selection gradients between the two treatment types using an ANCOVA to perform a regression for relative fitness on whitefly resistance, treatment, and their interaction.

### Greenhouse Experiment - Chlorophyll Content

To investigate the effects of dicamba on chlorophyll content in the greenhouse, we performed analysis of variance on the plants grown in the greenhouse. We fit the following simple linear model: y = u + treatment + e; where, y, the response variable, is chlorophyll average, u is the intercept or mean of the model, and e is the error term. In order to test for correlations between chlorophyll averages and whitefly abundance, we performed Pearson’s correlation test.

## Results

### Herbivory Response to Drift Exposed Host

We found no evidence that dicamba drift influenced the amount of chewing damage experienced by *A. theophrasti* (Fig 1A; STable 1) but we did find a significant increase in whitefly abundance on dicamba treated plants (F-value = 12.01, p-value < 0.01, Fig 1B; STable 1). This effect was also present in the second field season in which we included five more *A. theophrasti* populations (F-value = 5.30, p-value < 0.01, Fig 1C; STable 1). Both levels of dicamba drift -- 0.5% and 1% of the suggested field dose -- exhibited higher whitefly abundance than the controls. Plants exposed to 0.5% dicamba had significantly higher whitefly abundance than 1% dicamba treatments across both years (2018: t = 2.74, p < 0.01, Fig IB; 2019: t = 2.06, p = 0.04, Fig 1C).

**Figure 1.**
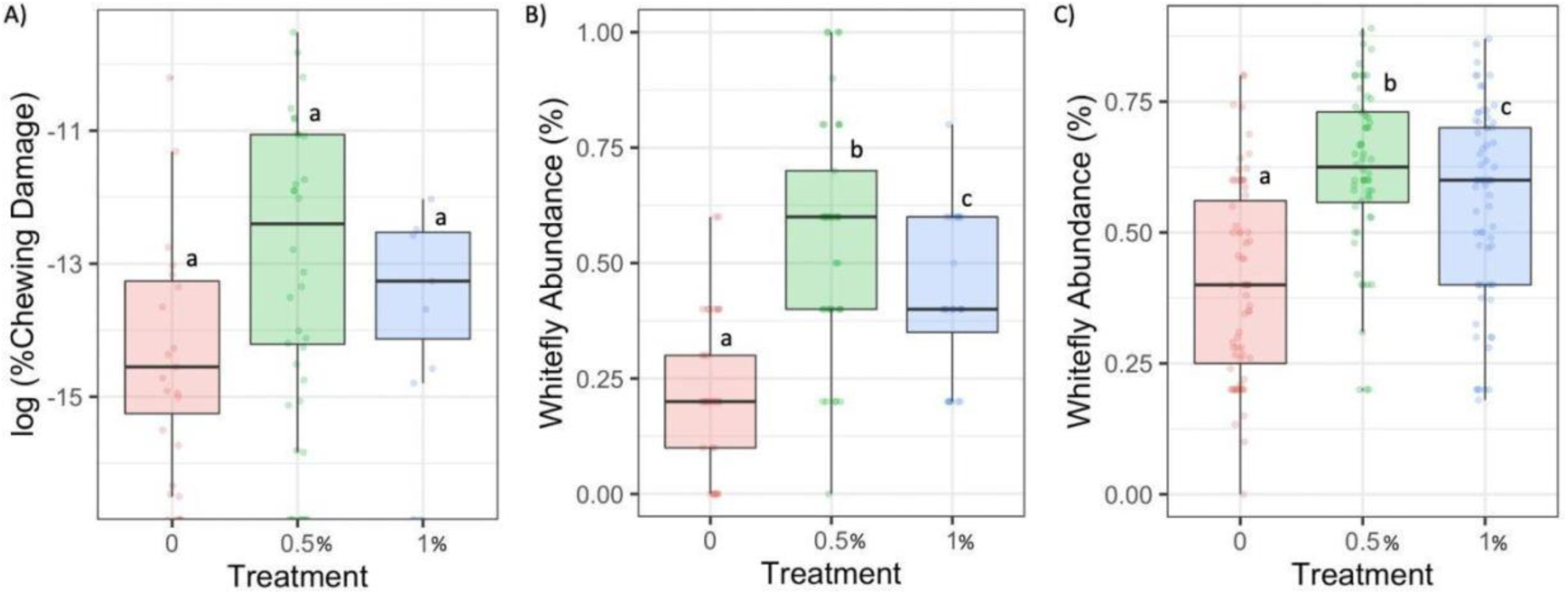
Chewing damage (A) and whitefly abundance (B and C) measurements in response to dicamba drift: 0% field dose (peach), 0.5% field dose (green), and 1% field dose (blue). A) Chewing herbivory damage summarized by dicamba treatment from the first field experiment in 2018. Damage was measured as area with chewing damage/total leaf surface area. B) Treatment effect on Whitefly abundance summarized by treatment in both the B) 2018 and C) 2019 field experiments. Whitefly abundance was measured as visual estimates of percent larvae area/total leaf surface area. Each graph illustrates median values and confidence intervals.

### Plastic Response to Herbicide Drift on Plants

We examined several plant traits in response to dicamba drift in order to determine how drift impacts velvetleaf growth and physiology, which may underscore herbivory responses. Across both experimental years, we found that leaf count increased following the application of dicamba drift (Treatment effect for 2018: F-value = 28.01, p-value < 0.01, Fig 2; Stable2; 2019: F-value = 5.11, p-value < 0.01, Fig 2; STable 3), whereas leaf width decreased significantly in 2018 but not 2019 (Treatment effect for 2018: F-value = 8.25, p-value < 0.01, Fig 2; STable 2; 2019: F-value = 0.79, p-value = 0.46, Fig 2; STable 3), which may be a result of significant population variation in 2019 for leaf width (χ^2^ = 5.11, p-value = 0.02, STable 3). Both plant height and flower count decreased as a result of dicamba drift in both experiments (Treatment effect: Height 2018: F-value = 220.56, p-value < 0.01; Flower Count 2018: F-value = 7.28, p-value < 0.01, Fig 2; STable 2; Treatment effect: Height 2019: F-value = 2.84, p-value = 0.06; Flower Count 2019: F-value = 6.92, p-value < 0.01, Fig 2, STable 3). Typically, plant leaf number was greater, and height and flower count were lower in the 1% dose of dicamba drift compared to the 0.5% dose (Fig 2, STable 2). We found evidence for block effects and block by treatment effects in the first field experiment across most phenotypic traits, but these effects were less evident in the second field experiment (STable 3). Finally, we found no evidence for maternal line or population effects associated with dicamba drift exposure, indicating that at least in this sample of 50 maternal lines, there was no evidence that the plastic response for height, leaf count, leaf width, and flower count varied genetically.

**Figure 2.**
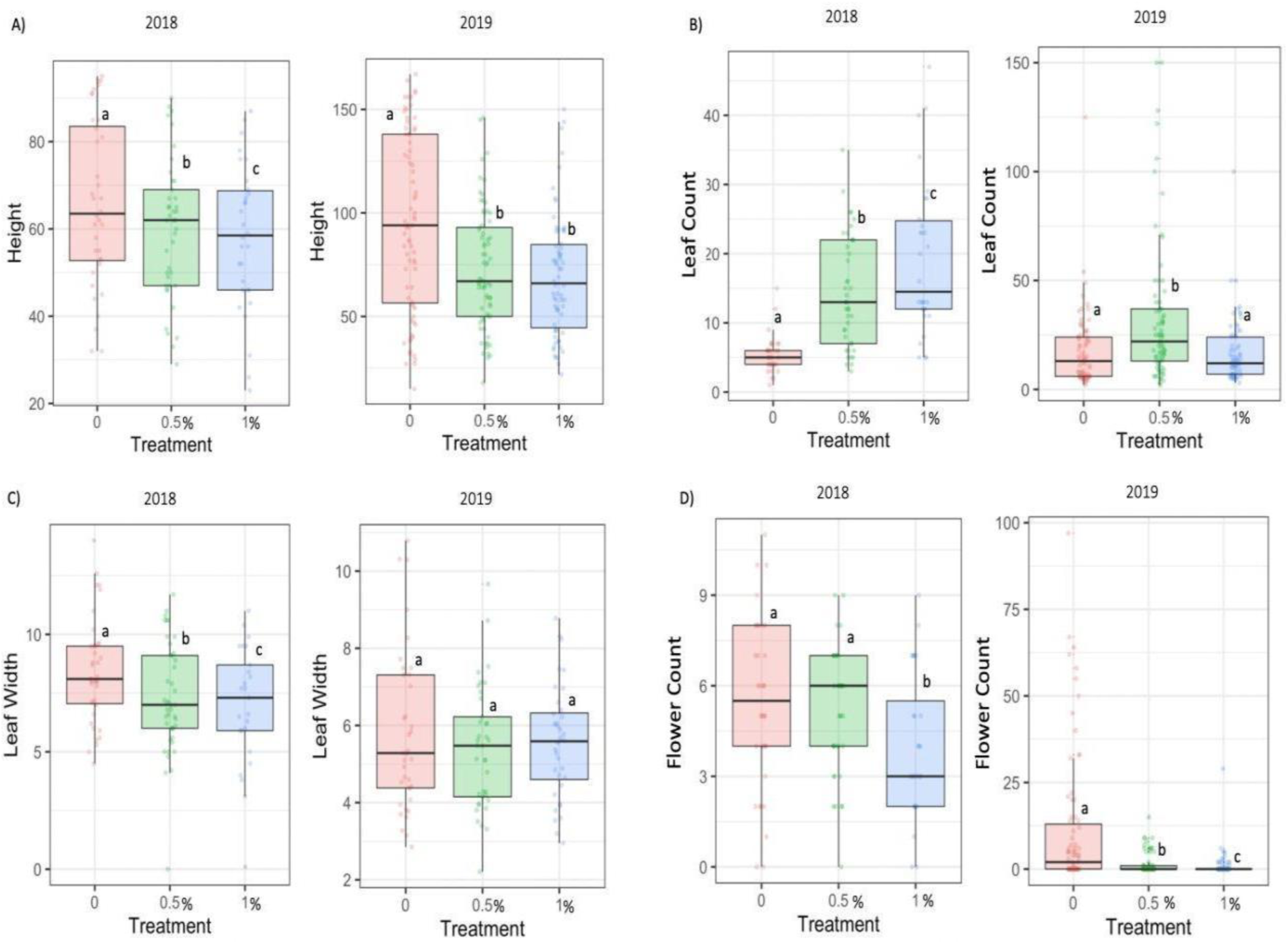
Plant size measurements in response to dicamba drift environments: 0% field dose (peach), 0.5% field dose (green), and 1% field dose (blue). Shows treatment effect for both years on velvetleaf traits: A) height B) leaf count C) leaf width and D) flower count. Each graph illustrates median values and confidence intervals.

Photosynthetic carbon dioxide response (*A*-*C*_i_, umol/m2), which was examined in the 2018 experiment, differed across treatments, indicating that dicamba drift significantly impacts photosynthetic carbon dioxide response overall (x^2^ = 37.605, p < 0.001; SFig 2;). Within drift treatments, we found evidence that time of leaf development significantly impacts A-Ci curves (0.5%: W = 696, p < 0.001 SFig 2; 1%: W = 360, p < 0.001). In general, leaves exposed directly to 0.5% of the field dose of dicamba exhibited higher A/Ci responses than their counterparts that developed after dicamba treatment (SFig 2). In contrast to the lower dosage, leaves treated directly with 1% of the field dose of dicamba exhibited lower A/Ci responses than their counterparts that developed after dicamba exposure (SFig 2). This result indicates high amounts of photosynthetic plasticity in response to dicamba, potentially associated with altered herbicide translocation and metabolism contributing to resistance to drift damage.

### Chlorophyll Content

In the greenhouse, we uncovered a significant treatment effect on chlorophyll levels (mg/cm^2^), where plants exposed to dicamba at drift levels had on average higher chlorophyll levels than the controls (F = 4.56, p = 0.03, Fig 3A). We also identified a significantly positive correlation between average chlorophyll values and whitefly abundance (r = 0.25, p < 0.001, Fig 3B), suggesting that chlorophyll content impacts whitefly host-selection on velvetleaf.

**Figure 3.**
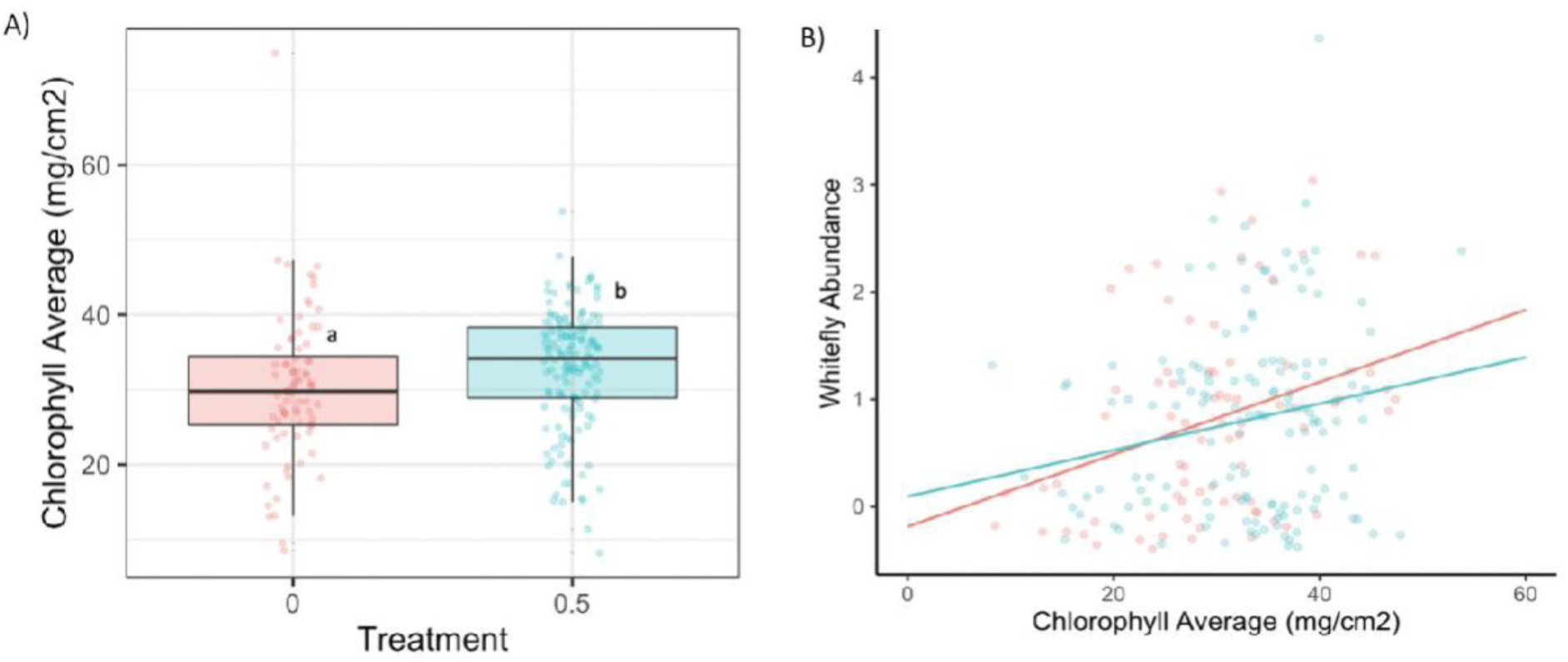
*Greenhouse Experiment* A) Average chlorophyll content per individual in response to dicamba application of 0.5% of the field dose (F = 4.56, p = 0.03). B) Relationship chlorophyll average and whitefly abundance estimated as Pearson product-moment correlation; Control treatment (peach) r = 0.34, p = 0.002; Drift treatment (blue) r = 0.21 p = 0.01. Each graph illustrates median values and confidence intervals.

### Genetic variation for whitefly resistance, herbicide resistance, and relative growth rate

We did not detect evidence of maternal line, maternal line by treatment, population, or population by treatment effects on whitefly resistance, herbicide resistance or relative growth rate in either year. (2018: STable 4, 2019: STable 5). This suggests there is not significant genetic variation in the populations we sampled from, likely due, at least in part, to the low number of maternal lines included in the study.

### Phenotypic correlations

We did not detect evidence of phenotypic correlations between any of the focal traits in 2018 (*i*.*e*. herbicide resistance, whitefly resistance, and relative growth rate), but in 2019 we found evidence of a moderately strong negative phenotypic correlation between herbicide resistance and whitefly resistance within drift environments (r = -0.32, p > 0.001). This suggests that both types of resistance may be indirectly impacting one another. We also found a negative correlation between whitefly resistance and relative growth rate in the presence of drift but not in its absence in 2019 (drift environment: r = -0.22, p = 0.02; control environment: r = -0.07, p = 0.58), indicating a tradeoff between growth and herbivory defense when exposed to herbicide drift.

### Phenotypic selection

In 2018, we did not identify evidence of linear selection on herbicide resistance nor whitefly resistance (herbicide resistance: β = -0.19, p = 0.57; whitefly resistance: β = 0.08, p = 0.30; STable 4) in the herbicide drift environment, though we did identify selection acting on relative growth rate (β = 1.71, p < 0.001; STable 4). We further detected marginal evidence of quadratic selection acting on whitefly resistance (γ = -0.19, p = 0.09, STable 4) and marginal correlative selection acting on whitefly resistance and relative growth rate (γ = 0.82, p = 0.07, STable 4). In 2019, we again found no evidence of linear selection in the herbicide drift environment and identified positive selection on relative growth rate (whitefly resistance, β = 0.002, p = 0.99; herbicide resistance, β = 0.05, p = 0.72; relative growth rate, β = 2.17, p < 0.001; STable 5). Patterns of correlative selection differed in 2019 compared to the 2018 experiment in that we detected marginal evidence of correlative selection acting on herbicide resistance and relative growth rate (γ = 0.66, p = 0.08, STable 5). Overall, though we found a significant negative phenotypic correlation between resistance to herbicide and whitefly herbivory, indicating a trade-off between the two types of resistance (at least in 2019) we found no evidence of correlative selection acting on the two traits in either year (2018: γ = 3.36, p = 0.14; STable 4; 2019: γ = 0.29, p = 0.46; STable 5).

## Discussion

### Ecological effects -- dicamba drift increases whitefly abundance and alters plant morphology and physiology

Our study provides the first evidence that dicamba drift can increase phloem feeding herbivore abundance in the field, a finding that should be of concern to agriculture more broadly given the negative effects of whiteflies on crops (*e*.*g*., viral transmission mediated by whitefly - Hogenhout 2008, Legg et al. 2014, Ning et al. 2015, Sundararaj et al. 2018, Moodley et al. 2019). In two separate field experiments, velvetleaf exposed to dicamba drift showed higher whitefly abundance in comparison to their control counterparts. These findings indicate dicamba drift can increase susceptibility to phloem feeding herbivorous insects such as whitefly, which is aligned with previous work showing increased abundance of English green aphids feeding on dicamba treated barley in a glasshouse experiment (Hintz 1971).

Additionally in the second year of our study, we found evidence of significant phenotypic tradeoffs between whitefly resistance and herbicide resistance as well as between whitefly resistance and relative growth rate in the presence of drift. Growth rate has long been linked to investment towards herbivory defense, suggesting that when resources are limited competition favors fast-growing plant species that allocate less towards herbivore defenses (Coley, Bryant, and Chapin 1985). In the literature, this is evident by positive correlations between growth rate and herbivory damage (Coley 1987, Cebrian and Durate 1994, Fine et al. 2006) and negative correlations between biomass accumulation and defense chemicals, such as salicylic acid (Meyer et al. 2007), a plant hormone specifically involved in defense against phloem-feeding silverleaf whiteflies (Kempema 2007). Such negative correlations suggest that pools of herbivore defenses can be depleted as strong growth occurs under certain environmental conditions. While our results reveal no correlation between growth rate and herbicide resistance, a significant correlation between growth rate and whitefly resistance when under drift induced stress supports the growth-defense tradeoff expectations of depleted reservoirs of herbivore defenses in the presence of high growth.

Because plant architecture is known to influence herbivore performance and abundance (Jaenike 1978, Haysom and Coulson 1998, Schlinkert et al. 2015), we assessed the morphological and physiological response of velvetleaf to dicamba drift. While our results indicate dicamba at drift levels stunts velvetleaf height, this synthetic auxin also increased the number of leaves produced, although they were typically smaller in size compared to non-sprayed control plants. Additionally, we examined the impact of dicamba on A/Ci response curves, as photosynthetic capabilities are linked to natural auxin presence (McAdam et al. 2007) and are a key driver for plant metabolism. The pattern of A/Ci response differed between drift treatments and time of leaf development, indicating a high amount of photosynthetic plasticity which varied over herbicide exposure and time. These findings align with a recent study that showed dicamba exposure initially decreased photosynthesis by 22% in *Palmer amaranth*, which then improved over time (Browne et al. 2020). Moreover, there was no evidence of maternal line nor population variation for morphological or physiological responses to herbicide drift, suggesting a high level of plasticity among all traits measured across individuals.

Natural auxins are involved in chlorophyll accumulation (Yuan et al. 2018). However, the addition of synthetic auxins, such as dicamba applied at 50% of the field dose (280g ae) causes greater than 70% reduction in chlorophyll content (Turgut 2007). Our results show that dicamba applied at drift rates (0.5% the field dose, i.e., 2.8g ae) can *increase* the average amount of chlorophyll in plants which could impact host-selection of whitefly. These findings are consistent with recent studies in which the greenhouse whitefly laid more eggs on leaves with higher chlorophyll and higher nitrogen (Park et al. 2009, Tsueda 2014) due to the highly linked nature of chlorophyll content and chloroplast thylakoids, which represent a large proportion of leaf nitrogen content (Evans 1983, Evans 1989), but conflicts with past work reporting whitefly attraction to yellow plants (Mound 1972, Van Lenterern and Noldus 1990). This suggests that color alone may not be the primary indicator for whitefly host-selection with every species. Although further studies on this dynamic are warranted, our results suggest that one mechanism behind the increase of whiteflies on dicamba-drifted plants is due to the auxin-induced increase in chlorophyll content of leaves.

### Evolutionary effects -- correlated evolution between resistance and relative growth rate

In addition to morphological and physiology responses, we also examined the potential that resistance to both dicamba drift and whitefly damage could evolve within these populations and investigated the potential for correlative selection between traits since we identified significant negative phenotypic correlations between herbicide resistance and whitefly resistance along with whitefly resistance and relative growth rate in 2019. In 2018, we detected marginal significance for correlative selection acting on whitefly resistance and relative growth rate. Correlative selection was also marginally detected in 2019 between relative growth rate and herbicide resistance in the presence of drift. While this could imply populations may evolve to an optimum fitness at intermediate values of relative growth rate, whitefly resistance and herbicide resistance, we did not uncover evidence of genetic variation within these populations. We further did not uncover significant maternal line variation for resistance traits, likely due to high amounts of plasticity within this species and/or an insufficient number of individuals per maternal line per treatment. Environment may also be a factor as previous studies in *Amaranthus hydrius* have shown that the fitness cost of herbicide resistance associated with increased herbivore susceptibility can be environmentally dependent (Gassmann 2005). Although we did not detect maternal line variation, more work here should be done investigating selection acting on the relationship between herbicide resistance, whitefly resistance, and relative growth rate with a greater number of maternal lines from more distant populations and environments under consideration.

### Conclusion

Herbicide drift from synthetic auxins can shift herbivorous insect host-selection at the agro-eco interface. Such modifications of herbivore behavior have the potential to impact plant community composition and nutrient dynamics (Schowalter 2006, Belovsky & Slade 2000) and may likewise, reciprocally influence insect herbivore populations. Increases in insect herbivore abundance may result in positive feedbacks where elevated consumption increases nutrient cycling and thus stimulates insect population growth, which can directly affect the abundance of other members of the community such predators and pollinators (Forkner & Hunter 2000, Hunter 2001, Ceulemans 2017).

Furthermore, our finding that dicamba drift significantly increases whitefly abundance on velvetleaf populations could mean dicamba-treated weeds act as a reservoir for whitefly populations, potentially leading to negative impacts on agricultural yields. If our results are applicable more broadly, plants exposed to dicamba drift in nature may be preferred host for whitefly populations. As agricultural pests that colonize more than 600 host plants globally (Byrne and Bellows 1991), whiteflies are known vectors for transmitting over 70% of the world’s plant viruses (Hogenhout 2008). Given the projected expanded use of dicamba, there is a clear and urgent need to examine to what extent community dynamics may shift as a result of dicamba use, especially for communities existing at the intersection of natural and managed vegetative system.

## Supplementary Tables and Figure

Treatment and Block impact on Herbivory

**STable 1.** Influence of herbicide treatment on chewing damage for 2018 and whitefly abundance for both years, analyzed using *F*-statistics values showing effects of treatment, block, treatment by block interactions, and likelihood ratio test statistics (x^2^) showing maternal line variation on herbivory measurements. In 2019 maternal lines are nested within populations. Significant values are expressed in boldface.

| Herbivory Response | Chewing Damage (2018) |  | Whitefly Abundance (2018) |  | Whitefly Abundance (2019) |  |
| --- | --- | --- | --- | --- | --- | --- |
| Fixed Effects: | *F* | P | *F* | P | *F* | P |
| Treatment (df = 2) | 0.906 | 0.410 | **12.014** | **< 0.001** | **5.300** | **0.006** |
| Block<br>2018 (df = 1)<br>2019 (df = 2) | 1.236 | 0.270 | 1.656 | 0.201 | 1.356 | 0.261 |
| Treatment X Block<br>2018 (df = 2)<br>2019 (df = 4) | 0.109 | 0.897 | 0.363 | 0.697 | 1.076 | 0.370 |
| Random Effects: | $\chi^2$ | P | $\chi^2$ | P | $\chi^2$ | P |
| Population (df = 1) |  |  |  |  | 0.000 | 1.000 |
| Maternal Line (df = 1) | 0.498 | 0.480 | 0.000 | 1.000 | 0.535 | 0.464 |
| Maternal Line X Treatment (df = 1) | 0.000 | 1.000 | 0.000 | 1.000 | 0.000 | 1.000 |

2018 Herbicide impacts on Velvetleaf Morphology

**STable 2.** 2018 influence of herbicide treatment on velvetleaf traits, analyzed using F-statistics values showing the effects of treatment, block, treatment by block interactions, and likelihood ratio test statistics **(χ2)** maternal line and maternal line by treatment interactions on variation of plant phenotypes. Significant values are expressed in boldface.

| 2018 |  |  |  |  |  |  |  |  |
| --- | --- | --- | --- | --- | --- | --- | --- | --- |
| Trait | Height |  | Leaf Count |  | Leaf Width |  | Flower Count |  |
| Fixed Effects: | *F* | P | *F* | P | *F* | P | *F* | P |
| Treatment<br>(df = 2) | **220.555** | **< 0.001** | **28.012** | **< 0.001** | **8.246** | **0.001** | **7.284** | **0.001** |
| Block<br>(df = 1) | 0.278 | 0.599 | **7.627** | **0.007** | **6.151** | **0.014** | **5.421** | **0.022** |
| Treatment X Block<br>(df = 2) | **60.804** | **< 0.001** | **15.714** | **< 0.001** | **3.554** | **0.032** | **4.287** | **0.016** |
| Random Effects: | $\chi^2$ | P | $\chi^2$ | P | $\chi^2$ | P | $\chi^2$ | P |
| Maternal Line<br>(df = 1) | 0.000 | 1.000 | 0.000 | 1.000 | 0.000 | 1.000 | 0.000 | 1.000 |
| Maternal Line<br>X Treatment<br>(df = 1) | 0.000 | 1.000 | 1.986 | 0.159 | 0.000 | 1.000 | 0.054 | 0.816 |

2019 Herbicide impacts on Velvetleaf Morphology

**STable 3.** Influence of herbicide treatment on velvetleaf traits, analyzed using F-statistics values showing the effects of treatment, block, treatment by block interactions, and likelihood ratio test statistics **(χ2)** showing population, maternal line, population by treatment interactions, and maternal line by treatment interactions on variation of plant phenotypes. Maternal lines were nested within populations. Significant values are expressed in boldface.

| 2019 |  |  |  |  |  |  |  |  |
| --- | --- | --- | --- | --- | --- | --- | --- | --- |
| Trait | Height |  | Leaf Count |  | Leaf Width |  | Flower Count |  |
| Fixed Effects: | *F* | P | *F* | P | *F* | P | *F* | P |
| Treatment<br>(df = 2) | 2.844 | 0.061 | **5.113** | **0.007** | 0.791 | 0.458 | **6.916** | **0.001** |
| Block<br>(df = 2) | 0.144 | 0.866 | 0.700 | 0.498 | 2.398 | 0.096 | 1.313 | 0.271 |
| Treatment X Block<br>(df = 4) | 0.551 | 0.699 | 0.506 | 1.697 | 1.763 | 0.188 | 0.470 | 0.758 |
| Random Effects: | $\chi^2$ | P | $\chi^2$ | P | $\chi^2$ | P | $\chi^2$ | P |
| Population<br>(df = 1) | 0.000 | 1.000 | 0.632 | 0.427 | 5.112 | 0.024 | 0.397 | 0.528 |
| Maternal Line<br>(df = 1) | 0.000 | 1.000 | 0.000 | 1.000 | 0.000 | 1.000 | 0.166 | 0.684 |
| Maternal Line<br>X Treatment<br>(df = 1) | 0.000 | 1.000 | 0.000 | 1.000 | 0.000 | 1.000 | 0.000 | 0.986 |

2018 Selection Analysis

**STable 4.**
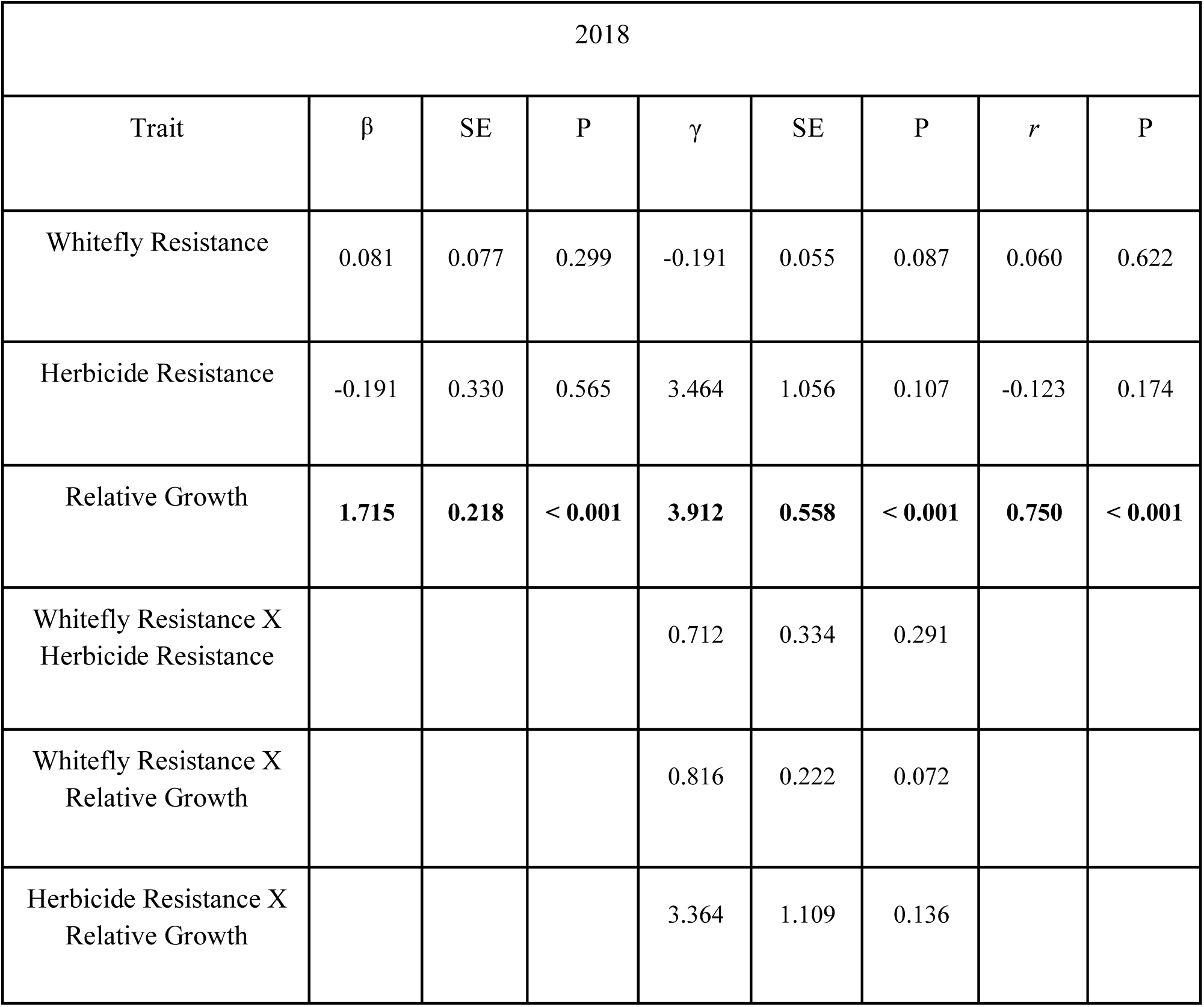
2018 Selection analysis showing direct selection on focal traits: whitefly resistance, herbicide resistance, and relative growth rate. Linear (β) (R^2^ = 0.510; p < 0.001) and quadratic (γ) (R^2^ = 0.609; p < 0.001) selection gradients with associated standard errors (SE) and *P*-values (*P*). The (r) column represents correlation coefficients between trait and fitness, estimated as Pearson product-moment correlations. Significant values are expressed in boldface.

| 2018 |  |  |  |  |  |  |  |  |
| --- | --- | --- | --- | --- | --- | --- | --- | --- |
| Trait | $\beta$ | SE | P | $\gamma$ | SE | P | $r$ | P |
| Whitefly Resistance | 0.081 | 0.077 | 0.299 | -0.191 | 0.055 | 0.087 | 0.060 | 0.622 |
| Herbicide Resistance | -0.191 | 0.330 | 0.565 | 3.464 | 1.056 | 0.107 | -0.123 | 0.174 |
| Relative Growth | **1.715** | **0.218** | **< 0.001** | **3.912** | **0.558** | **< 0.001** | **0.750** | **< 0.001** |
| Whitefly Resistance X<br>Herbicide Resistance |  |  |  | 0.712 | 0.334 | 0.291 |  |  |
| Whitefly Resistance X<br>Relative Growth |  |  |  | 0.816 | 0.222 | 0.072 |  |  |
| Herbicide Resistance X<br>Relative Growth |  |  |  | 3.364 | 1.109 | 0.136 |  |  |

2019 Selection Analysis

**STable 5.**
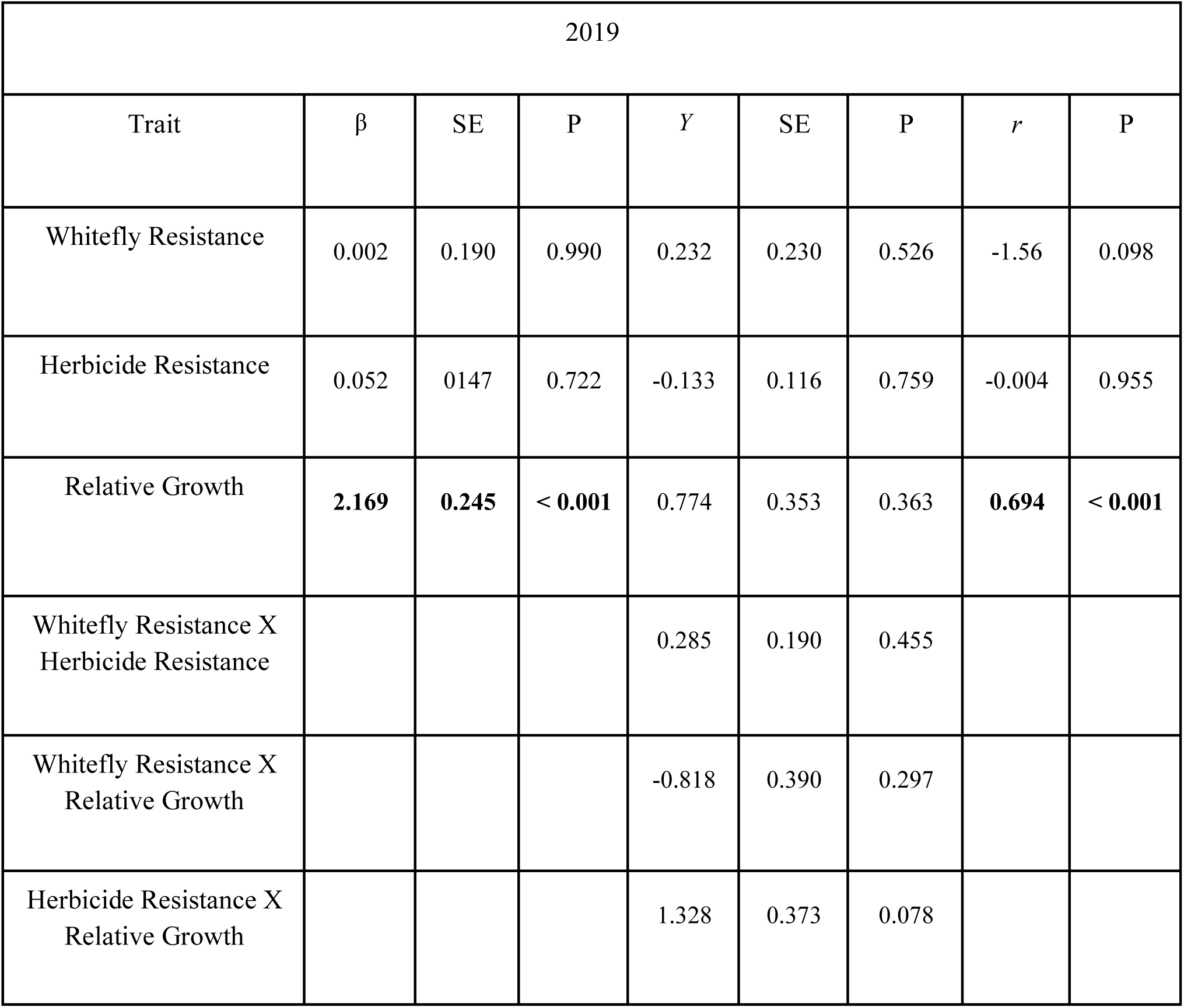
2019 Selection analysis showing direct selection on focal traits: whitefly resistance, herbicide resistance, and relative growth rate. Linear (β) (R^2^ = 0.453; p < 0.001) and quadratic (γ) (R^2^ = 0.466; p < 0.001) selection gradients, and total selection with associated standard errors (SE) and *P*-values (*P*). The (r) column represents correlation coefficients between trait and fitness, estimated as Pearson product-moment correlations. Significant values are expressed in boldface.

2018 Treatment and Block impact on Relative Growth Rate and Herbicide Damage

**STable 6.**
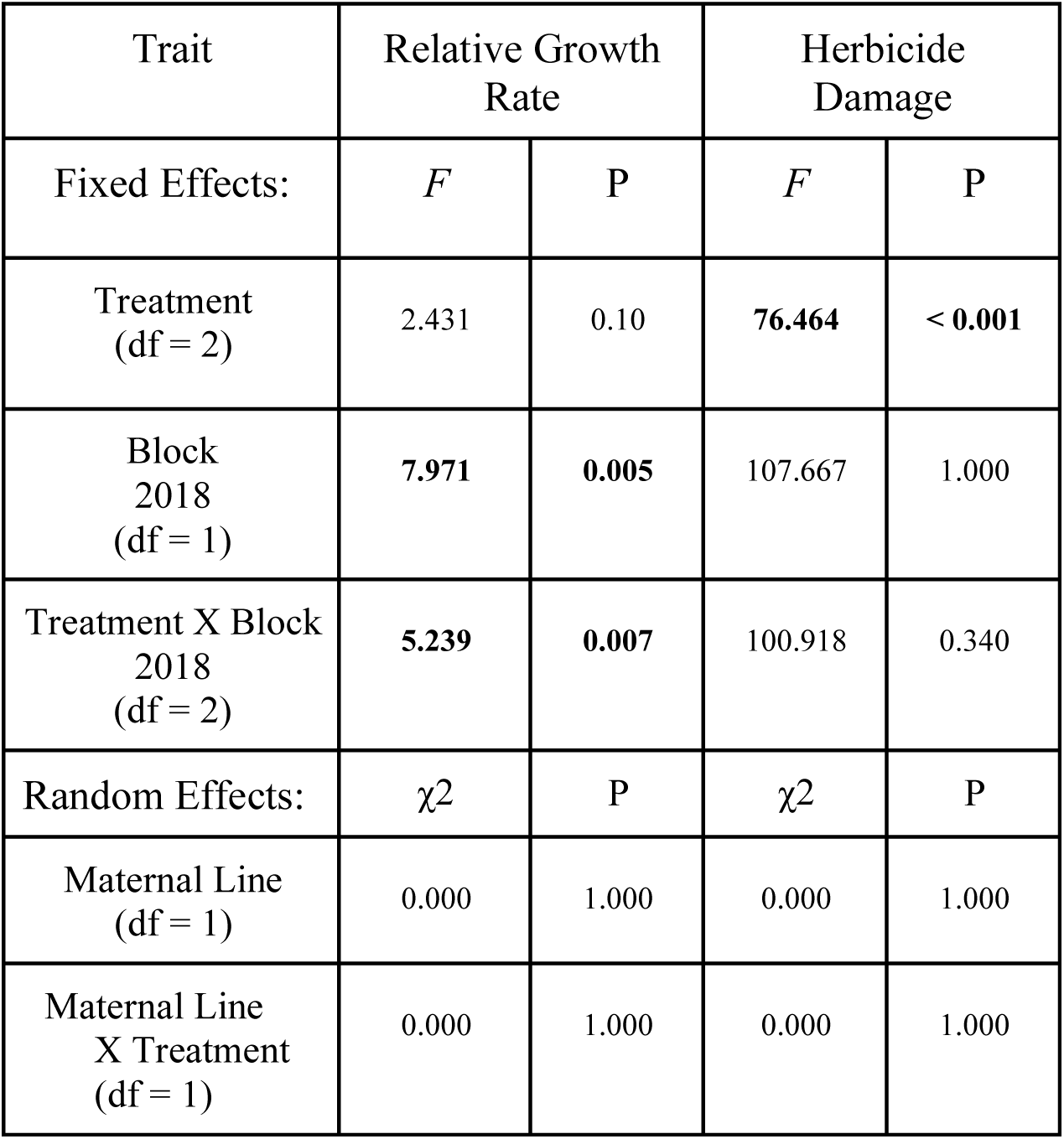
2018 Influence of treatment and block on relative growth, and herbicide damage, analyzed using F-statistics values showing the effects of treatment, block, treatment by block interactions, and likelihood ratio test statistics **(χ2)** showing, maternal line, and maternal line by treatment interactions on variation of plant phenotypes. Significant values are expressed in boldface.

2019 Treatment and Block impact on Relative Growth Rate and Herbicide Damage

**STable 7.** 2019 Influence of treatment and block on relative growth, and herbicide damage, analyzed using F-statistics values showing the effects of treatment, block, treatment by block interactions, and likelihood ratio test statistics **(χ2)** showing population, maternal line, population by treatment interactions, and maternal line by treatment interactions on variation of plant phenotypes. Significant values are expressed in boldface.

| Trait | Relative Growth Rate |  | Herbicide Damage |  |
| --- | --- | --- | --- | --- |
| Fixed Effects: | *F* | P | *F* | P |
| Treatment<br>(df = 2) | 0.031 | 0.969 | **46.730** | **< 0.001** |
| Block<br>(df = 2) | 0.323 | 0.724 | 0.006 | 0.994 |
| Treatment X Block<br>(df = 4) | 0.259 | 0.904 | **5.982** | **< 0.001** |
| Random Effects: | $\chi^2$ | P | $\chi^2$ | P |
| Population<br>(df = 1) | 0.000 | 1.000 | 0.190 | 0.663 |
| Population<br>X Treatment<br>(df = 1) | 0.479 | 0.488 | 2.300 | 0.129 |
| Maternal Line<br>(df = 1) | 0.000 | 1.000 | 0.000 | 1.000 |
| Maternal Line<br>X Treatment<br>(df = 1) | 0.000 | 1.000 | 0.000 | 1.000 |

**SFigure 1.**
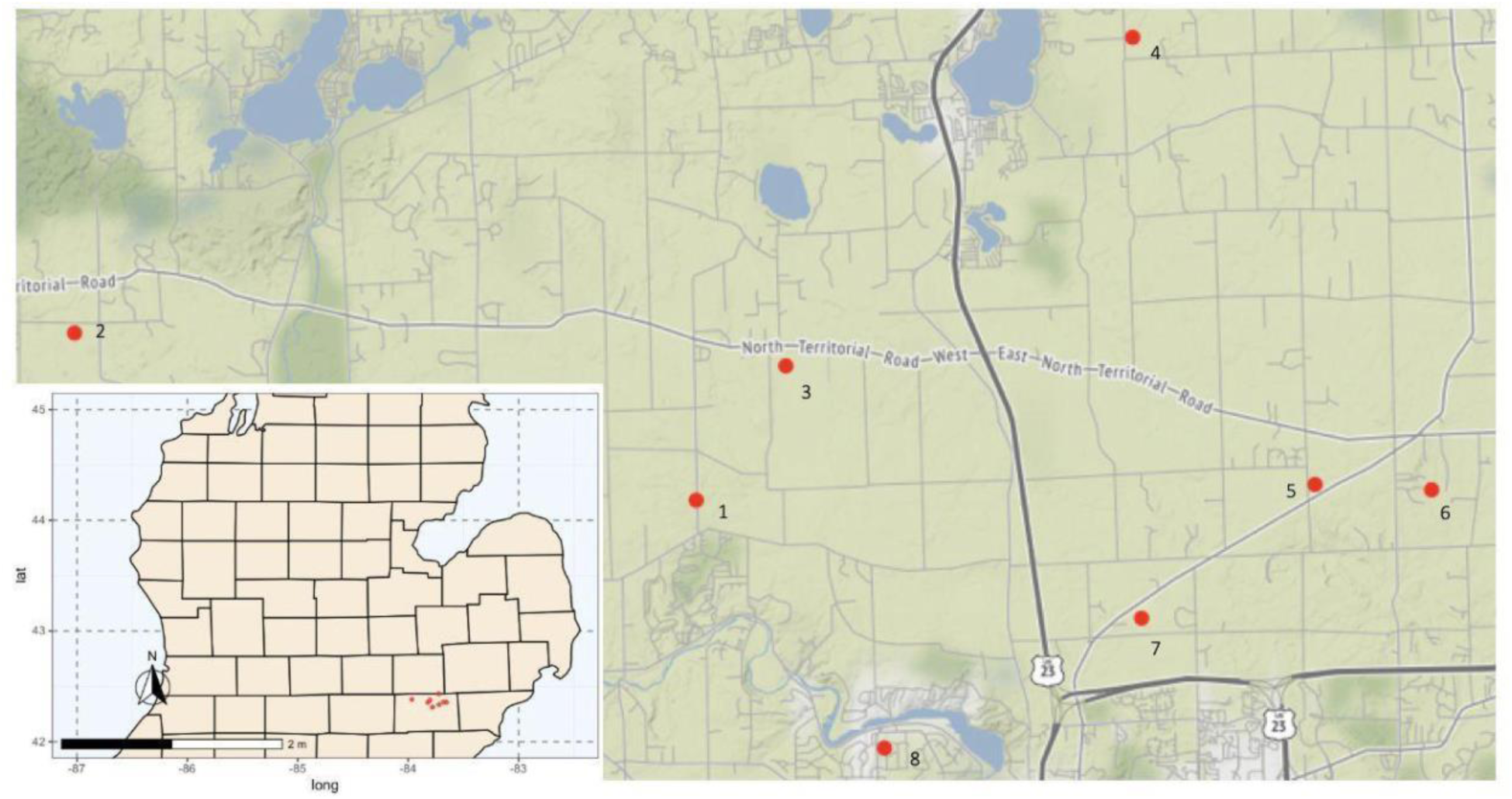
Locations of velvetleaf populations sampled and used for this study. In 2018, the field experiment was conducted with only population one, while in 2019 the sample size was increased to all eight populations. Populations 3, 4, 6, 7, and 8 were used in the 2021 greenhouse experiment.

**SFigure 2.**
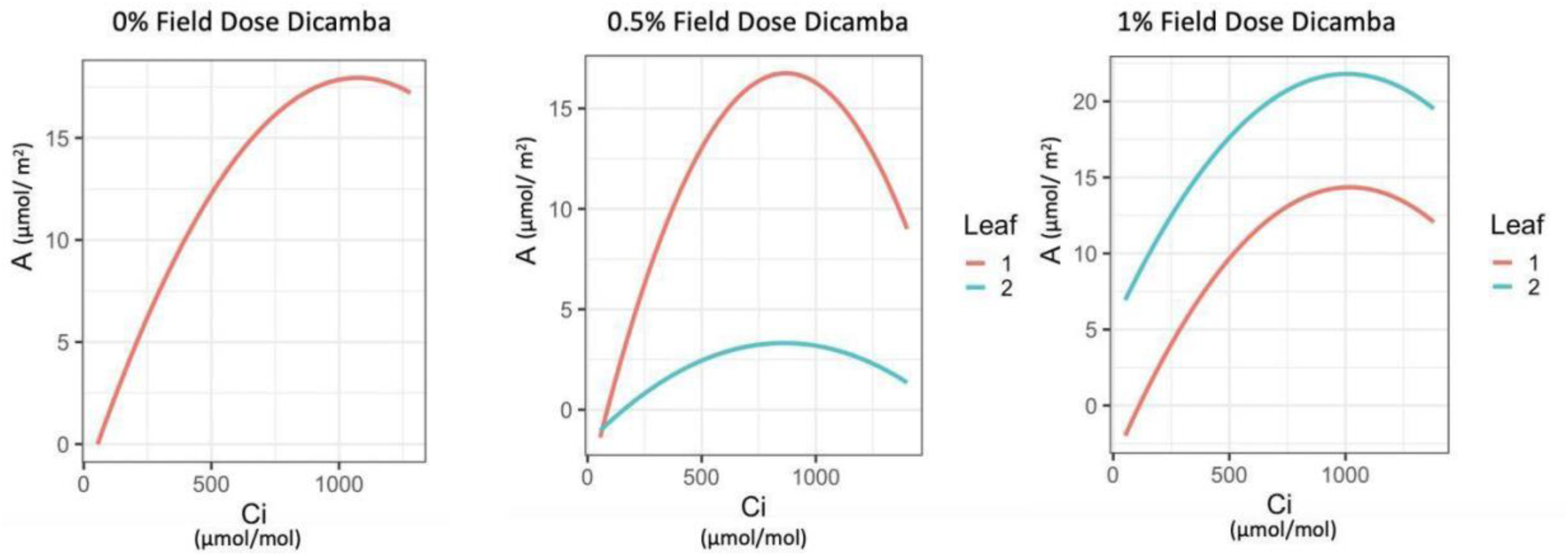
Photosynthetic carbon dioxide response curves by drift environment. A) *A*-*C*_i_ curves measured on leaves grown without drift exposure B) Comparison of *A*-*C*_i_ curves measured on leaves that developed before drift exposure (Leaf 1) and after drift exposure (Leaf 2) at 0.5% field dose. C) and at 1% field dose.

